# YOLito: A generalizable model for automated mosquito detection

**DOI:** 10.1101/2025.11.20.689454

**Authors:** Evyatar Sar-Shalom, Ziv Kassner, Arad Sarig, Clément Vinauger, Iliano Coutinho-Abreu, Merybeth F. Triana, Lucía I. Bouzada, R. Jason Pitts, Marcus C. Stensmyr, Omar S. Akbari, Philippos A. Papathanos, Jonathan D. Bohbot

## Abstract

Understanding mosquito behavior is key to advancing research in ecology, evolution, and disease control, yet most behavioral assays rely on human-dependent methods, such as real-time observation or manual frame-by-frame annotation, limiting throughput and reproducibility. We present YOLito, a domain-generalized AI model for automated mosquito detection and behavioral quantification. Built on the Ultralytics YOLO framework and enhanced with Slicing-Aided Hyper Inference (SAHI), YOLito accurately detects multiple mosquitoes across diverse backgrounds and imaging conditions. Trained on a globally assembled dataset of 38,547 annotated images from 35 experimental setups across six laboratories and three public datasets, YOLito achieved high performance on unseen data (precision = 0.95; recall = 0.91) and generalized across mosquito species (*Aedes, Anopheles, Culex*) and assay types, including blood-feeding, sugar-feeding, and oviposition. By automating behavioral scoring, YOLito transforms traditional assays into scalable and reproducible experimental platforms. The accompanying open-source toolkit enables high-throughput extraction of metrics such as visit frequency, duration, and distance traveled, providing a standardized and extensible framework that bridges computer vision and vector biology.

## Background

Observing animal behavior is fundamental to understanding species biology, ecology, and evolution^1,2^. Advances in digital imaging and high-resolution recording now allow precise capture of complex behaviors, but behavioral annotation remains a major bottleneck. Most analyses still rely on manual, frame-by-frame scoring, limiting throughput, reproducibility, and the scale at which behavior can be studied.

Automated annotation methods based on deep learning have begun to address this challenge. Frameworks such as DeepLabCut^3^ and DeepEthogram^4^ use convolutional neural networks (CNNs) to automate pose estimation and action classification. However, these approaches typically demand both extensive manual annotation and significant computational resources to achieve effective training, and retraining is typically necessary when experimental conditions change. Pretrained object detection frameworks such as Faster R-CNN^5^ and YOLO^6^ offer potential solutions but lack robustness in experimental variable settings, where lighting, background, and object size differ across biological scales^7-10^.

In mosquito behavioral research, these limitations are especially pronounced. Mosquitoes are small, fast-moving, and morphologically similar across species and sex, while experimental setups vary widely between laboratories. Existing object detection datasets for mosquitoes typically consist of static, high-resolution images of single or preserved specimens, which are unsuitable for dynamic behavioral analysis. Consequently, no current framework enables reliable, cross-laboratory quantification of mosquito behavior under diverse environmental and experimental conditions.

To overcome these challenges, we developed a domain-generalized AI model for automated mosquito detection and behavioral quantification named YOLito. Built on the Ultralytics YOLO framework and enhanced with Slicing Aided Hyper Inference (SAHI)^11^, YOLito achieves highly accurate detection of various sizes and resolutions of small, moving mosquitoes across complex and variable backgrounds (**Fig. 1**). The model was trained on a globally assembled dataset of 38,547 annotated images representing 35 experimental setups from six laboratories, supplemented by three public datasets. This diversity enables strong generalization across mosquito species (*Aedes, Anopheles, Culex*) and behavioral assays, including blood-feeding, sugar-feeding, and oviposition.

**Figure 1.**
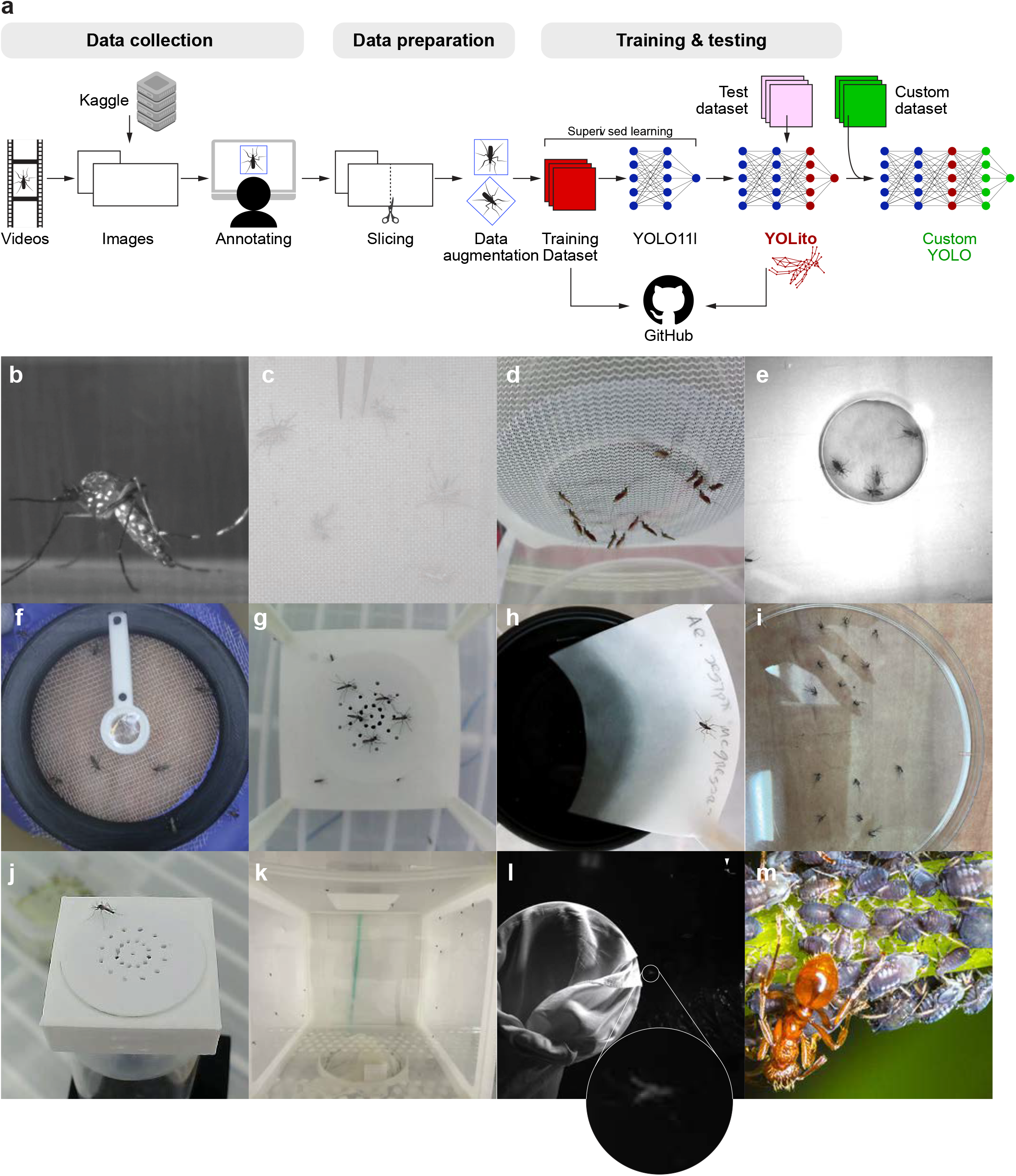
Overview of the YOLito model development pipeline. **a.** Videos recorded against varying backgrounds produced images of different sizes and visual characteristics. Mosquitoes were manually annotated, and the images were sliced using SAHI to preserve feature detail, then augmented using a range of techniques. The resulting dataset was used to train YOLO, producing the YOLito model. Both the training dataset and pretrained YOLito weights are available on GitHub. Because YOLito was trained to recognize general insect features, it can be further fine-tuned for detecting other insect species using additional custom data. Representative image types showing the variety of our mosquitoes’ dataset, including **b**. Macro image. **c**. High-resolution top and side view. **d**. Side view of blood-feeding. **e-i**. Bird’s-eye view of: **e**. blood-feeding, **f**-**h**. Landing, walking and egg-laying. **i**. Dead specimen. **j**. Diagonal view. **k**. Side view at flight. **l**. Grayscale flight **m**. background.

YOLito replaces manual annotation with automated, reproducible detection and quantification, transforming traditional behavioral assays into scalable and standardized experimental platforms. Trained to recognize general insect features, YOLito can serve as a pretrained model for a wide range of insect species, enabling rapid adaptation to new behavioral contexts. As a result, this approach represents a methodological advance that not only streamlines mosquito behavior analysis but also paves the way for automated, high-throughput behavioral assays in other insect species, including the black soldier fly, vinegar fly, honeybee, and Mediterranean fruit fly (medfly).

The accompanying open-source toolkit provides accessible post-detection analytics, enabling extraction of behavioral metrics such as visit frequency, duration, and distance traveled. This combination of domain-generalized learning and user-friendly analysis establishes a framework that is both user-accessible and extensible.

Together, YOLito and its companion training dataset bridge computer vision and vector biology, providing a unified tool for behavioral quantification across experimental and ecological contexts. Beyond mosquitoes, this approach offers a broadly applicable foundation for high-throughput behavioral research in other small-animal systems, accelerating discoveries in ecology, neuroethology, and disease-vector control.

## Methods for data collection

### Data collection

To develop a domain-generalized mosquito detector, we assembled a broadly representative dataset of 9,158 images sourced from 35 experimental setups contributed by six independent research groups, alongside publicly available images from Kaggle and recent studies^12,13^ (**Fig. 1**). The dataset includes a wide range of conditions relevant to mosquito research, such as top and side views (**Fig. 1b–c, j**), blood-feeding (**Fig. 1d-e**), oviposition (**Fig. 1h**), sugar foraging (**Fig. 1g**), flight (**Fig. 1k**), landing (**Fig. 1f**), and dead specimens (**Fig. 1i**). To improve specificity and reduce false positives, we also incorporated 367 non-mosquito insect images (**Fig. 1m**). The dataset spans substantial variation in brightness, contrast, resolution, and mosquito size (**Supplementary Fig. 1, Fig. 2**), ensuring that the trained model generalizes across diverse experimental designs and imaging conditions. By capturing this diversity,YOLito facilitates robust and accessible automated behavioral analysis across a range of vector biology applications. To support broad use, we released the annotated training dataset and YOLito on GitHub. YOLito can be further trained with custom annotations of the training dataset to detect additional features (*e*.*g*., body parts, behaviors, sex) or used with other frameworks like Faster R-CNN, Detectron, or newer YOLO versions.

**Figure 2.**
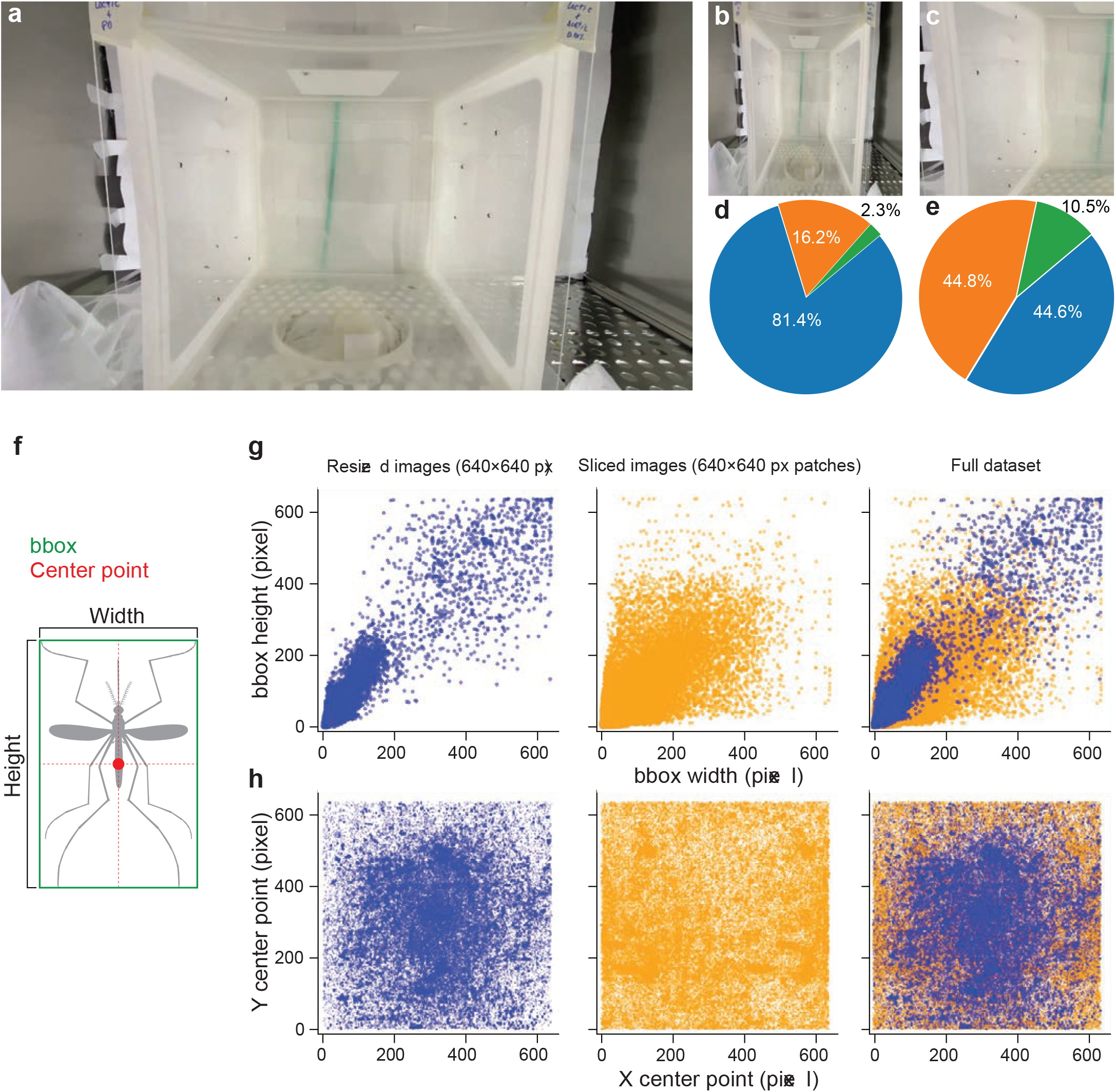
Balancing object size. **a-c.** A visual example of image transformations applied during training. **a**. An input image with small mosquito objects. **b**. Model’s perception of input images resized to 640×640 pixels. **c**. Model’s perception of sliced images as 640×640 pixel chunks (using SAHI). **d-e**. Distribution of bounding box sizes before (**d**) and after (**e**) slicing, according to MS COCO standards: small (<32×32 px), medium (32×32–96×96 px), and large (>96×96 px), labeled in blue, orange and green, respectively. **f**. Schematic representation of a bounding box (green) and the center point (red). **g**. Scatter plots of bounding box width vs.height for the resized, sliced, and full dataset. **h**. Spatial distribution of object coordinates (center X, Y) across the image frame for each dataset variant, showing object localization density.

### Balancing object size and enhancing feature visibility through slicing and augmentation

To enable consistent mosquito detection across diverse experimental conditions where object size and image resolution vary considerably, we standardized all training inputs to 640×640 pixels (**Fig. 1**). However, directly downscaling high-resolution images (**Fig. 2a**) compromises feature visibility by shrinking mosquito bounding boxes (bboxes), making it difficult for the model to learn relevant patterns (**Fig. 2b**). To mitigate this challenge, we implemented Slicing Aided Hyper Inference (SAHI), which divides large images into smaller, non-overlapping 640×640 tiles while preserving original object size and detail (**Fig. 2c**). Prior to slicing, 81.4% of objects were categorized as small, 16.2% medium, and 2.3% large (n = 101,856), according to MS COCO size definitions^14^ (**Fig. 2d**). Post-slicing, this shifted to 44.8% small, 44.6% medium, and 10.5% large (n = 139,080), resulting in a total of 240,936 annotated objects across the dataset (**Fig. 2e**). This approach yielded a 3.5-fold increase in average bbox area, from 23.38 (±45.54) × 30.56 (±50.54) pixels to 48.48 (±56.82) × 52.02 (±54.51) pixels, providing the model with larger object sizes (**Fig. 2g**). Additionally, slicing diversified bbox center points within the frame (**Fig. 2h**), reinforcing the model’s ability to detect mosquitoes across the entire image space. Slicing also produced a more balanced object-size distribution. To further improve generalization, we applied data augmentation, including HSV color shifts, horizontal and vertical flipping, scaling, mosaic augmentation, and random cropping (YOLO settings in **Extended Data 1**). These techniques increased dataset diversity, enabling the model to better handle variation in mosquito appearance and experimental conditions. The SAHI slicing tool was integrated into the YOLito codebase, allowing users to apply image slicing during both training and inference to enhance performance in high-resolution and small-object detection scenarios.

## Model training, validation, and benchmarking

### Training framework and optimization

To build YOLito, we selected the YOLOv11l architecture for its strong performance in detecting both small and large objects. Training was conducted using the Ultralytics ecosystem, which offers streamlined implementation, integrated support for image slicing, and robust documentation. The full dataset comprised 9,158 original mosquito and non-mosquito images, which expanded to 38,547 (30,840 and 7,707 split into training and cross-validation datasets, respectively) through SAHI preprocessing (**Supplementary Table 1, Fig. 2**). An additional 53 annotated images from unseen experiments served as a test dataset to evaluate the model’s generalization (**Supplementary Table 2**).

The model was trained for 800 epochs using a batch size of 32 and a fixed input resolution of 640×640 pixels. We applied Ultralytics’ default data augmentation techniques (**Fig. 1,3a**). To optimize learning, we used a warm-up phase with a constant learning rate of 0.01 for the first three epochs, followed by a 1% per-epoch decay, facilitating smooth convergence.

Loss optimization targeted three components:

- Box Loss (CIoU): Penalizes spatial mismatches between predicted and ground-truth bounding boxes.
- Classification Loss (Binary Cross Entropy): Improved mosquito classification accuracy.
- Distribution Focal Loss (DFL): Enhanced localization by modeling each coordinate as a discrete probability distribution.

The loss curves exhibited a sharp initial decline, reflecting effective early learning, and continued to decrease steadily across epochs without signs of overfitting (**Fig. 3b**). Performance metrics improved in parallel, reaching a mAP@0.5 of 0.94, mAP@0.5:0.95 of 0.67, precision of 0.94, and recall of 0.88 (**Fig. 4c–e**), indicating high detection accuracy and generalization potential.

**Figure 3.**
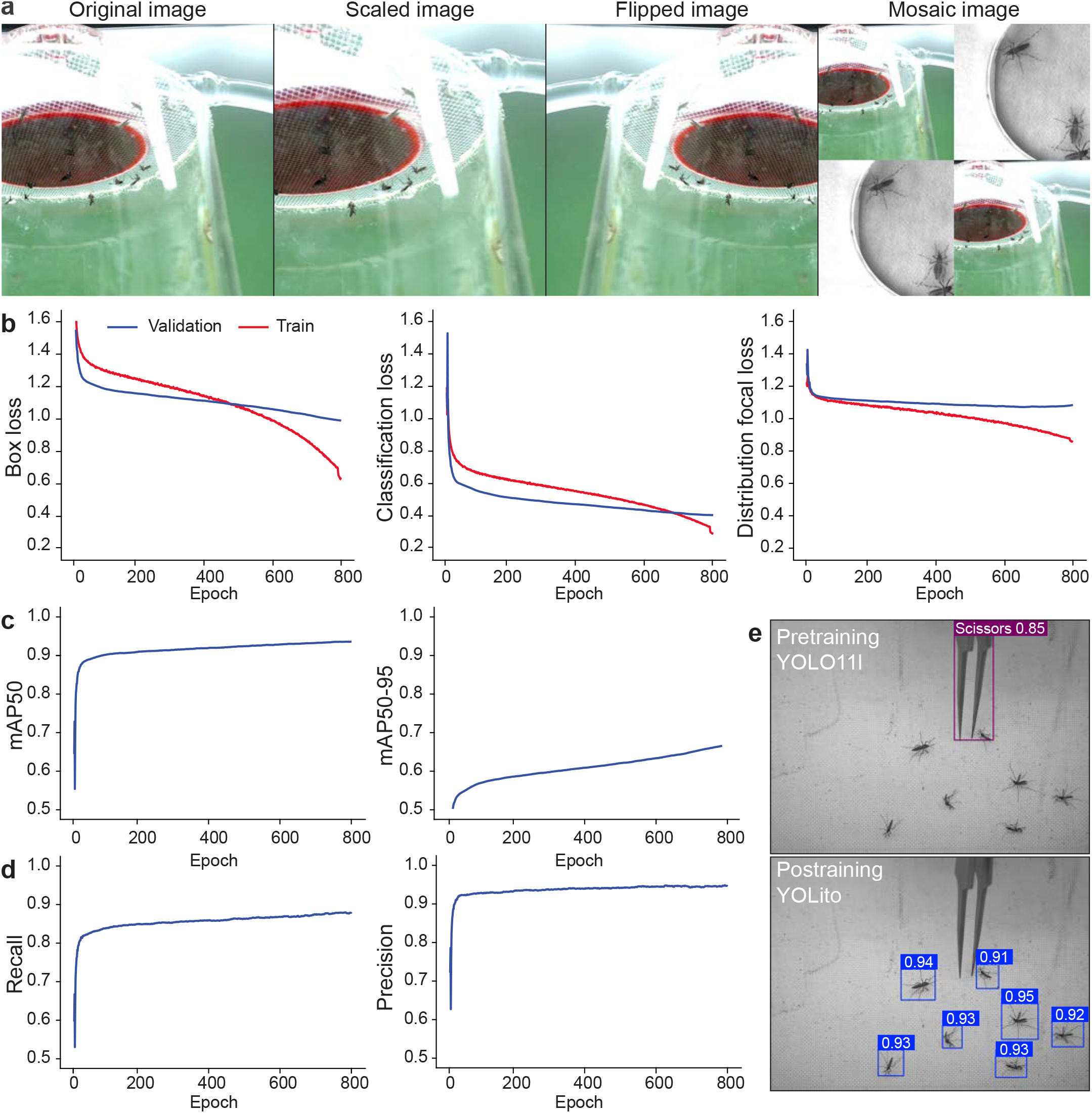
Training progress and validation. **a.** Representative data augmentations applied during training, including scaling, horizontal flipping, and mosaic. **b**. Training (red) and validation (blue) loss curves across 800 epochs for bounding box regression (Box), classification (Cls), and distribution focal loss (DFL). **c-d**. Model performance on the validation dataset during training, showing mean average precision at IoU threshold 0.5 (mAP50), and mAP averaged over IoU thresholds from 0.5 to 0.95 (mAP50–95) (c), recall, and precision (d). The model exhibits convergence and stable performance. **e**. Example detections before training (pretrained YOLO11l) and after fine-tuning (YOLito), illustrating transformation towards a domain-generalized mosquito detector.

**Figure 4.**
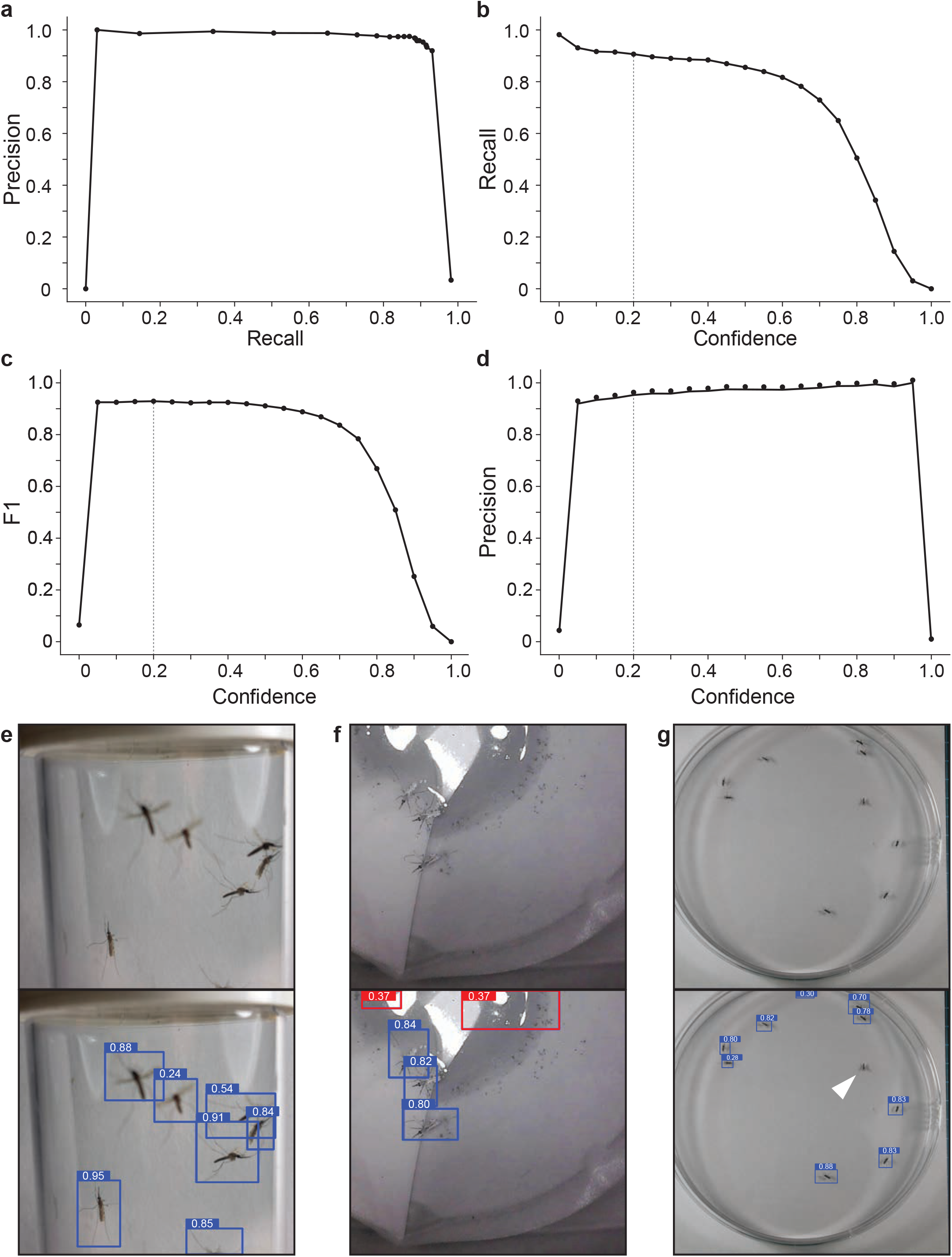
Generalization performance. **a.** Precision–recall curve across confidence thresholds. **b**. Recall as a function of confidence threshold. **c**. F1 score across confidence levels.**d**. Precision as a function of confidence threshold. (**e**–**g**) Prediction results in three representative experimental setups: **e**. *An. gambiae* in flight; **f**. *An. gambiae* during oviposition, illustrating a false positive detection labeled in red; **g**. *Ae. aegypti* in a Petri dish, illustrating a false negative (white triangle). The top row shows original images, the bottom row shows YOLito predictions with bounding boxes and associated confidence scores.

### Cross-validation performance

We continuously monitored performance on the cross-validation dataset using recall, precision, mAP@0.5, and mAP@0.5:0.95. These metrics comprehensively capture detection accuracy and localization precision:

- Recall: Ratio of correctly predicted mosquitoes to total ground-truth instances.
- Precision: Ratio of correct predictions to all predicted instances.
- mAP@0.5: Proportion of detections with ≥50% Intersection-over-Union (IoU) overlap.
- mAP@0.5:0.95: Stricter measure averaging AP across IoUs from 0.5 to 0.95 in 0.05 increments.

Because YOLito targets a single class (mosquito), mean average precision (mAP) is equivalent to AP. Continuous performance gains across all metrics confirm the model’s ability to distinguish mosquitoes from background noise and non-target organisms.

### Benchmarking and domain-generalization evaluation

To assess generalization, we benchmarked YOLito on a fully independent test dataset (**Fig. 1**) of 53 annotated images spanning four mosquito species (*An. gambiae, An. stephensi, Ae. aegypti*, and *Cx. quinquefasciatus*, **Supplementary Table 2**) and multiple behavioral contexts (*e*.*g*., flight, blood-feeding, sugar-feeding, oviposition, dead specimens^15^). These images were unseen during training or validation, ensuring unbiased evaluation.

Model performance was again evaluated using precision, recall, and F1 score across confidence thresholds from 0 to 1 in 0.05 increments (**Fig. 4a-d**). Optimal detection performance was achieved at a confidence threshold of 0.2 and a non-maximum suppression (NMS) threshold of 0.35, yielding a recall of 0.91 and a precision of 0.95. For comparison, the pretrained YOLO11l model evaluated on the same independent benchmark dataset reached a precision of0.47 and a recall of 0.33.

Three representative case studies illustrated the model’s behavior in real-world scenarios:

- Accurate detections (**Fig. 4e**)
- False positives (**Fig. 4f**)
- False negatives (**Fig. 4g**)

### Computational resources

All model training, inference, and evaluation were performed on a dedicated workstation equipped with an NVIDIA GeForce RTX 4090 GPU (24 GB VRAM), an Intel Core i9-13900K CPU, and 64 GB RAM, running Ubuntu 22.04.2 LTS. GPU acceleration used CUDA 12.6 with NVIDIA driver 560.94.

### Software

All analyses were conducted in Python 3.10.13 within a Conda environment on Ubuntu 22.04.2 LTS. Model training and inference were implemented using the Ultralytics YOLO framework together with SAHI for image slicing. Data processing and visualization relied on standard scientific Python libraries, including NumPy, pandas, OpenCV, Matplotlib, and Plotnine. All custom code for dataset preparation, model training, and evaluation is available at GitHub (URL).

## Demonstration of high-throughput behavioral profiling

Despite its broad applicability, YOLito may show reduced accuracy on grayscale recordings, likely due to the limited representation of grayscale data in the training set. To test its performance with grayscale images, we validated YOLito’s performance under naturalistic and variable lighting conditions. We conducted egg-laying experiments under controlled 12:12 light:dark cycles using a 3D-printed recording platform (**Fig. 5a**). Twenty post-mated adult female *Ae. albopictus* (Foshan-Pavia, FPA) and *Ae. aegypti* (Liverpool) were blood-fed under insectary conditions (26°C, 80% RH) and transferred after 3 days to 20 cm^3^ observation cages containing a central oviposition cup lined with Whatman filter paper and filled with distilled water. The same experimental procedure was followed for both species, with *Ae. albopictus* used at 6 days post-emergence and *Ae. aegypti* at 13 days post-emergence. Behavior was continuously recorded for 24 hours using an infrared-capable Arducam 1080p Day & Night Vision USB camera mounted above the oviposition cup across three biological replicates. This enabled non-invasive, high-resolution video capture across both light and dark periods. For analysis, we extracted a 6-hour time window, 3 hours before and 3 hours after lights-off, and processed all footage using the YOLito inference engine.

**Figure 5.**
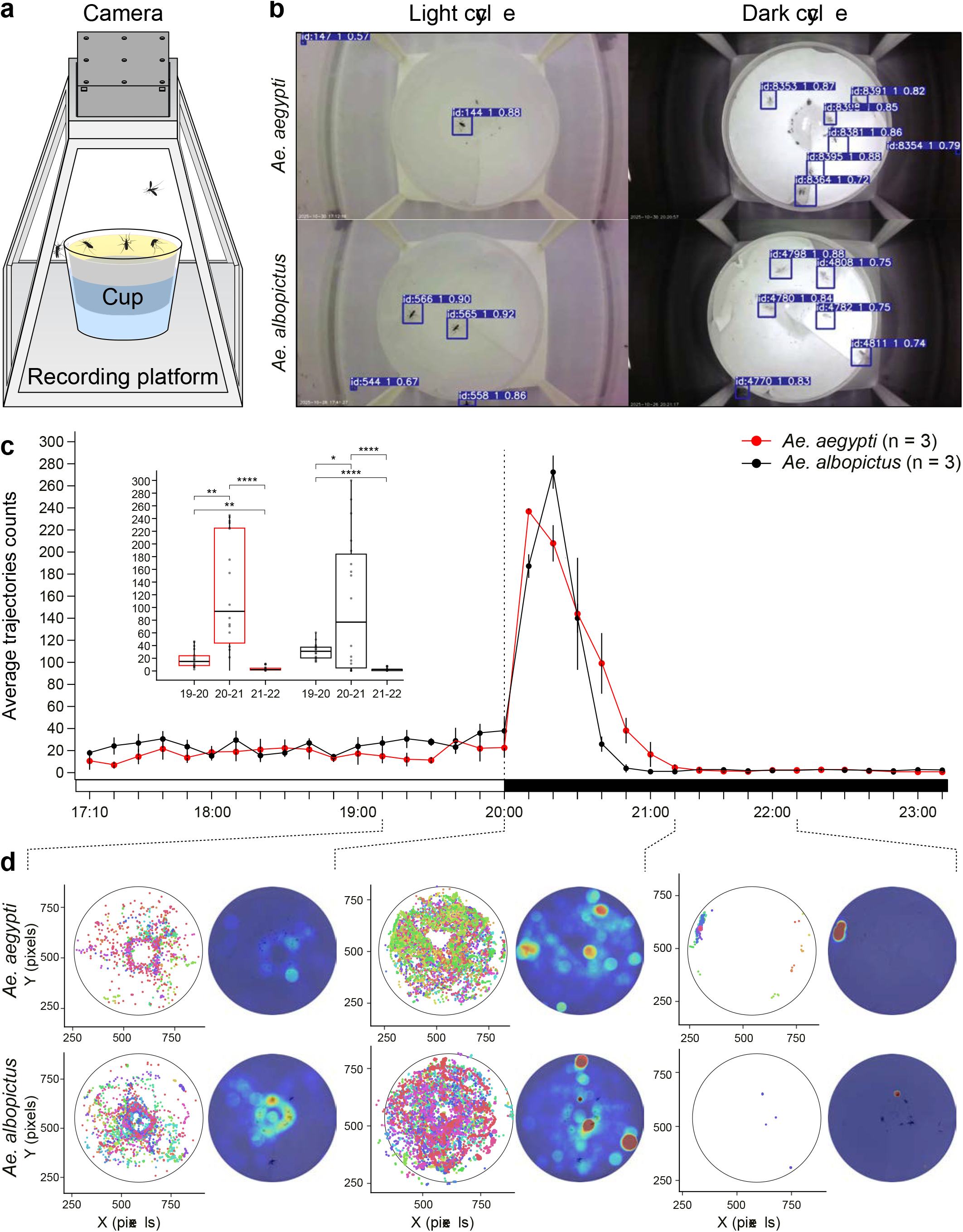
Detection of oviposition activity during light/dark cycles. **a.** Schematic of the oviposition recording setup. The ROI is the Whatman paper (yellow). **b**. Representative detections of visiting *Ae. aegypti* and *Ae. albopictus* females on the ROI during light and dark cycles. **c**. Temporal dynamics of mosquito trajectories visiting the ROI. The time between 17:10 and 23:00 is shown. The inset compares the number of trajectories at the peak activity level, one hour preceding, and following it. **d**. Representative coordinate distributions and activity heatmaps of visiting mosquitoes.

YOLito successfully detected mosquitoes during both the light and dark phases (**Fig. 5b**), enabling precise quantification of temporal visit dynamics of the oviposition cup. Both species exhibited low (basal) activity levels prior to the onset of darkness. Within minutes after the lights turned off, we observed a marked increase in visit frequency, peaking within the first hour and then tapering off for the remainder of the dark cycle (**Fig. 5c**). A generalized linear mixed-effects model (GLMM) confirmed that visit rates across the three temporal segments were significantly different within each species (**Fig. 5c** inset, **Supplementary Table 3**). No statistical difference was found between species in the same temporal segment (**Supplementary Table 4**).

Oviposition behavior was further visualized using the integrated post-detection analysis toolkit, which supports trajectory plots, spatial heatmaps, and temporal behavioral summaries (**Fig. 5d**). The toolkit includes the following modular functions:

- plotxy: Visualizes spatial distribution by plotting X-Y coordinates of tracked individuals across frames, revealing trajectories, clustering, and arena occupancy.
- heatmap: Generates overlay heatmaps with circular intensity effects centered on detections, highlighting preferred zones or avoided areas.
- visits: Computes visit frequency per interval based on unique track_id counts (Visits = count(unique(track_id))/interval), reflecting interaction rate with the region of interest (ROI).

All data, trained weights, and code for the inference and analysis pipeline are publicly available via the YOLitko GitHub repository, enabling broad accessibility and reproducibility across insect behavioral research. This framework transforms oviposition assays^16^ into scalable, automated, and quantitative platforms for high-throughput behavioral phenotyping in mosquitoes and other small insects.

## Avenues for future insect detection models

As a domain-generalized mosquito detection model, YOLito captures insect features that are likely shared across species. This makes it a strong candidate for use as a generalizable pretrained insect detector, which can be further customized through fine-tuning with custom datasets (**Fig. 1**). To evaluate this potential, we tested YOLito on images with varying sharpness from four non-mosquito insect species: the Mediterranean fruit fly, black soldier fly, honeybee, and vinegar fly. YOLito successfully detected all four species, albeit with varying confidence levels (**Fig. 6**). In general, precision exceeded recall, suggesting that while YOLito is sensitive to insect-like features, it exhibits a higher rate of false negatives, which will be reduced with targeted fine-tuning. Validation metrics ranked detection performance in descending order as: vinegar fly > black soldier fly > honeybee > Mediterranean fruit fly. Despite variation in insect size, morphology, and image sharpness, we acknowledge that this evaluation is limited to a single experimental setup with a uniform background. Future efforts to improve generalizability should include diverse imaging conditions, backgrounds, and viewpoints, as well as species-specific fine-tuning. Given its optimization for insect-relevant visual features, YOLito provides a robust foundation for transfer learning and can be more efficiently adapted to other insect species compared to training a model de novo (**Fig. 1**).

**Figure 6.**
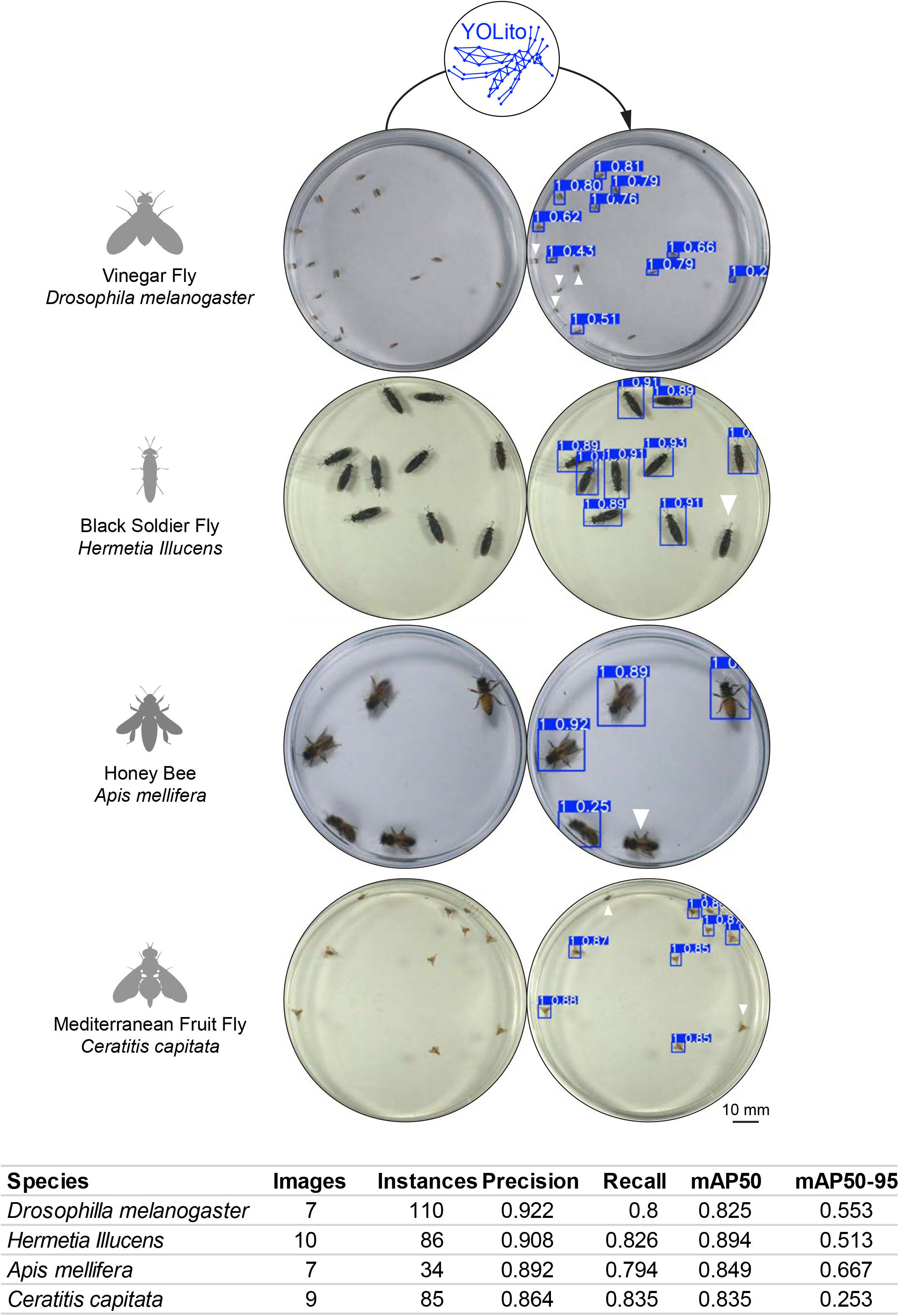
YOLito is a pretrained insect detection model. Detection of four insect species, including the Mediterranean fruit fly *Ceratitis capitata* (Diptera), the honeybee *Apis mellifera* (Hymenoptera), the vinegar fly *Drosophila melanogaster* (Diptera), and the black soldier fly *Hermetia illucens* (Diptera). Blue bounding boxes indicate positive detections, while false negatives are marked with solid white triangles. The accompanying Supplementary Table summarizes detection performance metrics across species.S

## Conclusions

YOLito introduces a scalable and domain-generalized framework for automated mosquito behavior analysis, replacing manual annotation with high-throughput, reproducible detection. Trained on a diverse dataset spanning four mosquito species, postures, and behavioral contexts, YOLito accurately detects mosquitoes across various lighting, resolutions, and experimental setups. By integrating image slicing (SAHI) and YOLO-based inference, it achieves high precision (0.95) and recall (0.91), even for small or partially occluded individuals. The open-source dataset, pretrained model, and modular analysis toolkit enable researchers to extract biologically meaningful metrics such as visit frequency, duration, locomotion, and space use with minimal coding expertise. Critically, YOLito was trained to recognize general insect features, making it a practical pre-trained model for broader entomological applications, including species like black soldier flies, Mediterranean fruit flies, and honeybees. By bridging computer vision with behavioral science, YOLito standardizes experimental workflows and supports the development of real-time, closed-loop, or large-scale assays. This methodological shift offers a powerful foundation for accelerating research in ecology, neuroethology, and disease-vector biology.

## Data availability

All data, weights, and analysis toolkit are available in our GitHub repository. https://github.com/WildMosquit0/YOLito

## Acknowledgments

This study was supported by ISF 719/21 awarded to J.B. This work was also supported by research grants from the Gates Foundation (INV-004363 and INV-075374 to PAP), the Israel Science Foundation (2388/19 to PAP), the Israel Ministry of Science & Technology (3-16795and 3-17985 to PAP) and the German-Israeli Middle East Project Cooperation of the German Research Foundation (1833/7-1 to PAP). The generation of behavioral data was partially supported by the National Institute of Allergy and Infectious Diseases of the National Institutes of Health under Award Number R01AI155785 (to C.V.) and the USDA National Institute of Food and Agriculture, Hatch Research project VA-160212 (to C.V.). O.S.A. and I.C. were supported by NIH grants (R01AI190001-01, RO1AI148300-05, and RO1AI175152-03). We would also like to acknowledge the NIH for funding support: This research was funded by the National Institutes of Health, National Institute of Allergy and Infectious Diseases, award number 1R01AI148300-01A1 to R.J.P.We thank Maree Kadee, Osnat Malka, Gleb Ens, Dor Peretz, Neta Keren, Yuri Vainer, and Ido Apel for providing insect specimens used for photography. The authors used an AI language model (ChatGPT, OpenAI) for editorial assistance. All scientific content and interpretations were developed and validated by the authors.

## Contributions

E.S.S. and J.D.B. conceived the project. E.S.S. developed YOLito and conducted the experiments. E.S.S., J.D.B. and Z.K. wrote the manuscript. Z.K. provided technical support on all aspects of YOLito’s development. A.S., P.A.P., C.V., I.C.A., and O.S.A. contributed to analysis design and data collection. M.F.T. and M.C.S., L.I.B. and R.J.P. contributed to data collection. All authors reviewed and approved the final version of the manuscript.

## Ethics declarations

O.S.A. is a founder of Agragene, Inc., Synvect, Inc., and Cloak with equity interest. The terms of this arrangement have been reviewed and approved by the University of California, San Diego in accordance with its conflict-of-interest policies. All other authors declare no competing interests.

## Supplementary Tables

**Supplementary Table 1. The full dataset consists of four components**: (1) images collected from six laboratories; (2) non-insect background images obtained from [Kaggle](https://www.kaggle.com/datasets/tarundalal/dangerous-insects-dataset); (3) random mosquito images from non-laboratory [Kaggle](https://www.kaggle.com/datasets/iloncka/mosquito-dataset-46-3139-segmentation-select-70) dataset; and (4) an open-source dataset from [Janson](https://www.frontiersin.org/journals/bioengineering-and-biotechnology/articles/10.3389/fbioe.2023.1103748/full?ref=toest.bg) et al. (2023).

**Supplementary Table 2. Test dataset**. Image dataset used to evaluate the YOLito model performance.

**Supplementary Table 3. GLMM analysis within species**. Pairwise contrasts between temporal segments within species.

**Supplementary Table 4. GLMM analysis between species**. Pairwise contrasts within temporal segments between species.

**Supplementary figure 1. Image features distribution across the dataset (Before slicing)**.

Histograms show the frequency of **a**. brightness, **b**. aspect ratio, **c**. contrast, **d**. width, **e**. height,**f**. entropy, **g**. sharpness, and **h**. saturation values calculated from all images. **i**. Correlation matrix depicting pairwise relationships between extracted image features.

**Extended data 1. YOLO settings for training**.

