## Supplementary figures and images for "YOLito: A generalizable model for automated mosquito detection"

### Supplementary Figure 1

#### Supplementary Figure 1

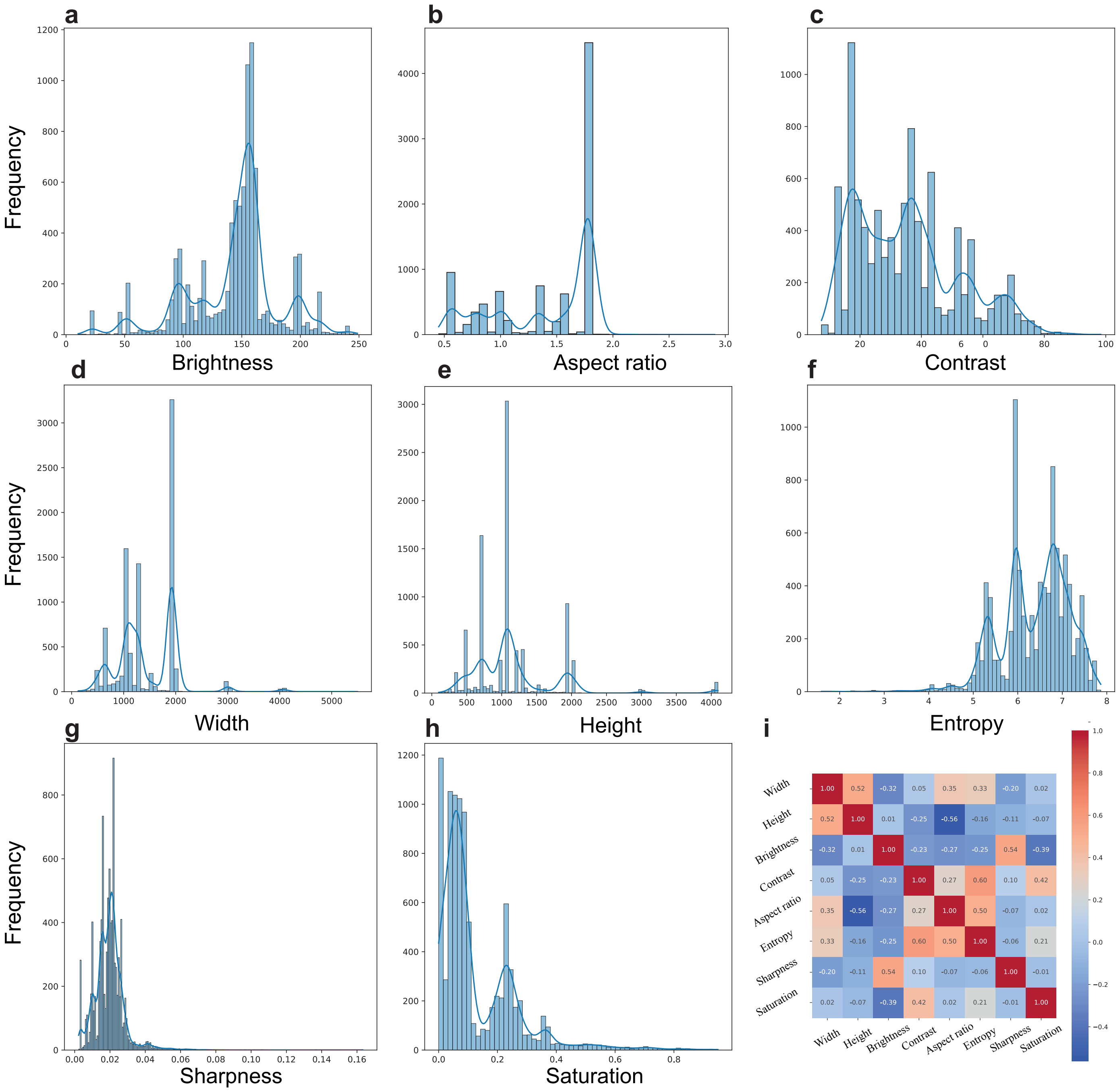
